# Spatial Needleman-Wunsch: A Deterministic Dynamic Programming Framework for Molecular Docking

**DOI:** 10.1101/2025.07.24.666489

**Authors:** Joshua Daniel Curry

**Affiliations:** Independent Researcher

**Keywords:** molecular docking, dynamic programming, voxelization, deterministic algorithms, multi-objective optimization, interpretable AI

## Abstract

Most molecular docking tools today rely on stochastic sampling or heuristic optimization. They can be effective, but reproducibility is often hit or miss, and there’s no guarantee the best binding pose has been found.

This paper introduces Spatial Needleman-Wunsch, a deterministic dynamic programming framework adapted from sequence alignment and applied to three-dimensional molecular structures. Ligands and protein cavities are voxelized into grids; chemical interactions are scored with compatibility matrices inspired by BLOSUM and PAM; and optimal alignments are computed exhaustively.

The result is a docking method that’s reproducible, interpretable, and mathematically grounded. The framework supports Boltzmann ensemble analysis, Pareto trade-off modeling, conformational sampling, and adaptive scoring. Proof-of-concept tests on synthetic systems and real targets like HIV protease show perfect reproducibility and meaningful chemical discrimination.

This approach eliminates random variability while making interaction scoring fully transparent. It opens the door to principled docking exploration, something essential in drug discovery and structural biology.

## 1 Introduction

Most molecular docking tools today embrace probabilistic search strategies. They rely on random sampling, heuristics, or genetic algorithms to approximate binding poses. While these techniques have enabled many breakthroughs, they often produce variable results, obscure the logic behind predictions, and offer no guarantee that the best solution has been found.

The field of bioinformatics, by contrast, is dominated by deterministic algorithms. The Needleman-Wunsch algorithm (4), for example, guarantees an optimal global alignment between two sequences by exhaustively evaluating possible matches using dynamic programming. Its success lies in combining thoroughness with efficiency through memoization, producing consistent and explainable results.

Current state-of-the-art docking tools, including AutoDock Vina (1), Glide (2), and GOLD (3), employ stochastic optimization strategies such as Monte Carlo sampling, genetic algorithms, and simulated annealing to explore the vast conformational space of protein-ligand interactions.

Despite their widespread adoption and practical utility, these methods face several persistent challenges:

1. **Stochastic variability**: Results can vary between runs due to random sampling, complicating reproducibility and systematic analysis
2. **No optimality guarantees**: Heuristic search methods may converge to local minima, potentially missing globally optimal binding configurations
3. **Limited interpretability**: Complex scoring functions and search algorithms can obscure the chemical rationale underlying predicted interactions

### 1.1 Motivation and Scope

This work sets out to ask: what if molecular docking worked more like sequence alignment? In bioinformatics, tools like Needleman-Wunsch offer guaranteed optimal matches between sequences. Could we bring that same clarity to spatial recognition in 3D chemistry?

By converting ligands and protein cavities into voxel grids, each with encoded chemical properties, we show that it’s possible to apply deterministic dynamic programming to molecular docking. The goal is to replace black-box heuristics with transparent, reproducible scoring.

This approach provides:

- Reproducibility (same input, same output)
- Scoring transparency via decomposable functions
- Flexibility for shape, charge, and accessibility comparisons
- Extensions for conformational freedom and multiple objective trade-offs

### 1.2 Relationship to Prior Work

While voxel-based representations have been explored in molecular modeling (7; 8), their application within a deterministic dynamic programming framework for docking represents a novel approach. Our method conceptually bridges sequence alignment algorithms with spatial optimization, creating a new paradigm for molecular recognition that prioritizes transparency and reproducibility.

## 2 Methods

### 2.1 Theoretical Foundation

The Needleman-Wunsch algorithm computes globally optimal alignments between biological sequences using dynamic programming. Its recurrence relation compares match/mismatch scores and gap penalties to build an optimal alignment path between two sequences. We adapt this concept to molecular docking by applying the same principles to three-dimensional molecular structures.

To do this, we discretize both ligands and protein cavities into voxel grids. Each voxel encodes chemical properties, allowing us to evaluate voxel-voxel interactions using a compatibility matrix, similar in spirit to amino acid substitution matrices.

The spatial alignment objective becomes:

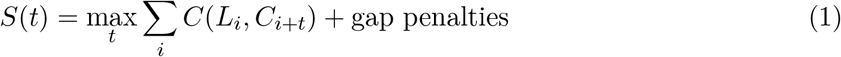

Where:

- *S*(*t*) is the alignment score for translation vector *t*
- *L*_*i*_ is the chemical type of voxel *i* in the ligand
- *C*_*i*+*t*_ is the corresponding voxel in the cavity after translation
- *C*(*a, b*) is the compatibility score between voxel types *a* and *b*

This formulation guarantees optimal spatial alignment under the defined scoring scheme, without relying on random sampling.

### 2.2 Voxel Representation

#### 2.2.1 Grid Generation

Molecules are converted into voxel grids, typically with a resolution of 0.5-1.0 Å per side. Each voxel represents a cubic unit of space and encodes three key properties:

- **Chemical type** (categorical): hydrophobic, polar, positively charged, negatively charged, aromatic, or empty
- **Accessibility** (continuous): a score between 0 and 1 representing solvent exposure
- **Flexibility** (continuous): reflects atomic mobility, often derived from B-factors

#### 2.2.2 Property Assignment

For protein cavities:

- **Proximity analysis** determines distances to nearby heavy atoms
- **Chemical environment** accounts for local residues and functional groups
- **Geometric constraints** define cavity shape and accessibility

For ligands:

- Each atom is mapped to its closest voxel center
- Chemical types are assigned using atomic properties, bonding patterns, and pharmacophoric features (e.g., hydrogen bond donors, aromatic rings)

### 2.3 Chemical Compatibility Matrix

To evaluate interactions, we define a compatibility matrix that scores voxel-voxel pairings. This matrix is inspired by physical chemistry and follows principles such as:

The matrix is extensible. Future versions may incorporate halogen bonding, metal coordination, or cation-*π* interactions.

### 2.4 Core Spatial Alignment Algorithm

The docking algorithm exhaustively tests all possible translations of the ligand within the cavity’s voxel grid, scoring each configuration using the compatibility matrix:

#### Algorithm 1

Spatial Alignment Algorithm

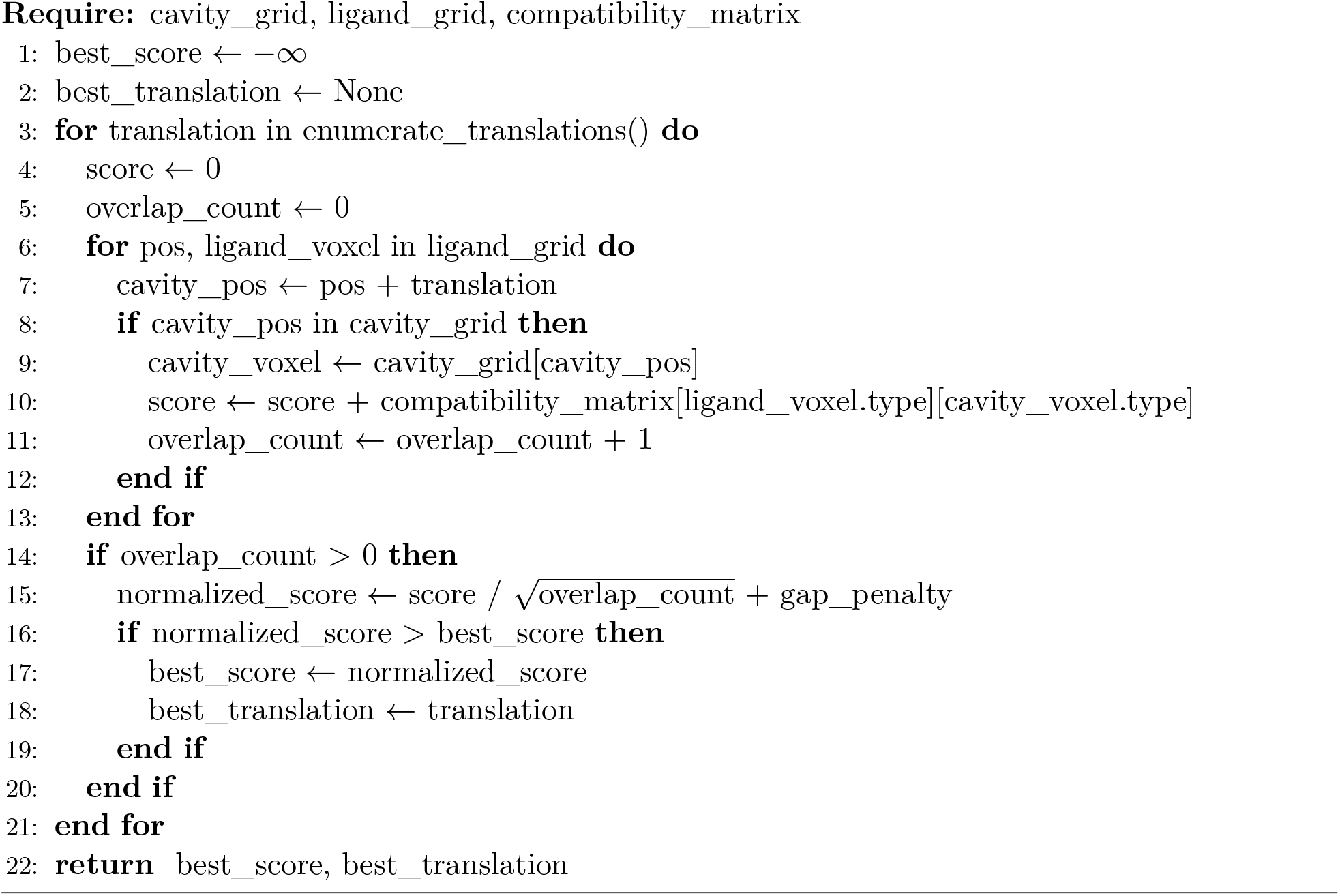

This search is guaranteed to find the optimal spatial alignment under the scoring model.

#### Computational Complexity

If the cavity has *N* ^3^ voxels and the ligand has *M*, the algorithm scales as 𝒪(*N* ^3^*M*). While computationally intensive, this remains tractable for most small-to-medium systems and could benefit from GPU parallelization.

### 2.5 Software Implementation

The Spatial Needleman-Wunsch framework is written in Python, with modular components for alignment, scoring, and visualization. The package structure includes:

- `core.py:` Main SpatialDocking and Voxel classes
- `alignment.py:` Translation scoring and dynamic programming core
- `scoring.py:` Compatibility matrix and evaluation logic
- `visualization.py:` 3D rendering utilities

Optional modules:

- `boltzmann.py:` Boltzmann ensemble analysis
- `pareto.py:` Multi-objective optimization
- `conformers.py:` Torsional sampling
- `adaptive.py:` Scoring refinement

#### 2.5.1 Core Classes

The framework centers on two primary classes:

##### Voxel Dataclass

Represents individual 3D grid points with chemical properties:

**Listing 1:**
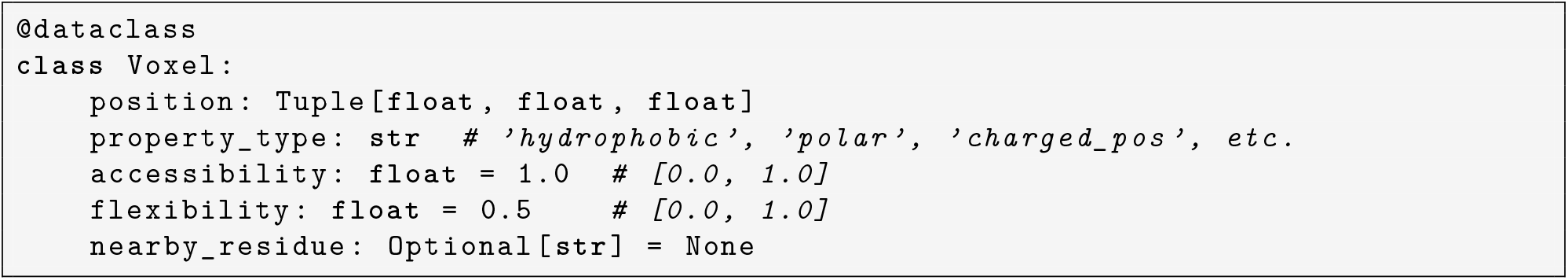
Voxel representation

##### SpatialDocking Class

Orchestrates the complete docking workflow:

**Listing 2:**
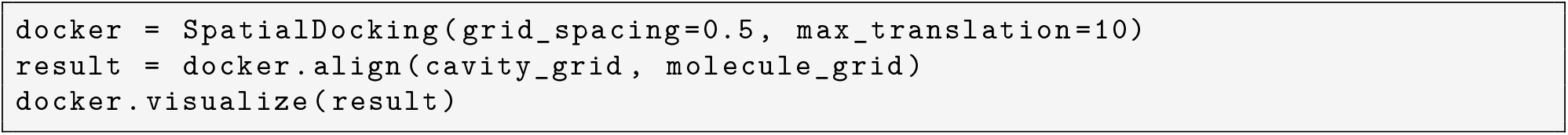
Basic usage example

#### 2.5.2 Result Object Structure

Each docking run returns a structured result object that contains:

- Optimal score and translation vector
- Input ligand and cavity grids
- Compatibility matrix used
- Full metadata for reproducibility

### 2.6 Framework Extensions

#### 2.6.1 Boltzmann Ensemble Analysis

Instead of reporting just the best pose, we compute a Boltzmann-weighted ensemble of all valid translations:

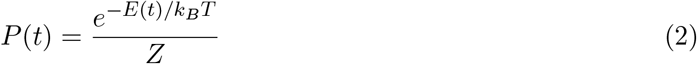

Where *E*(*t*) = −*S*(*t*) is the effective energy of pose *t*, and *Z* is the partition function. This allows ensemble-averaged analysis and entropy estimation.

#### 2.6.2 Pareto Frontier Optimization

Docking often requires balancing multiple criteria. The framework supports multi-objective optimization using:

**Listing 3:**
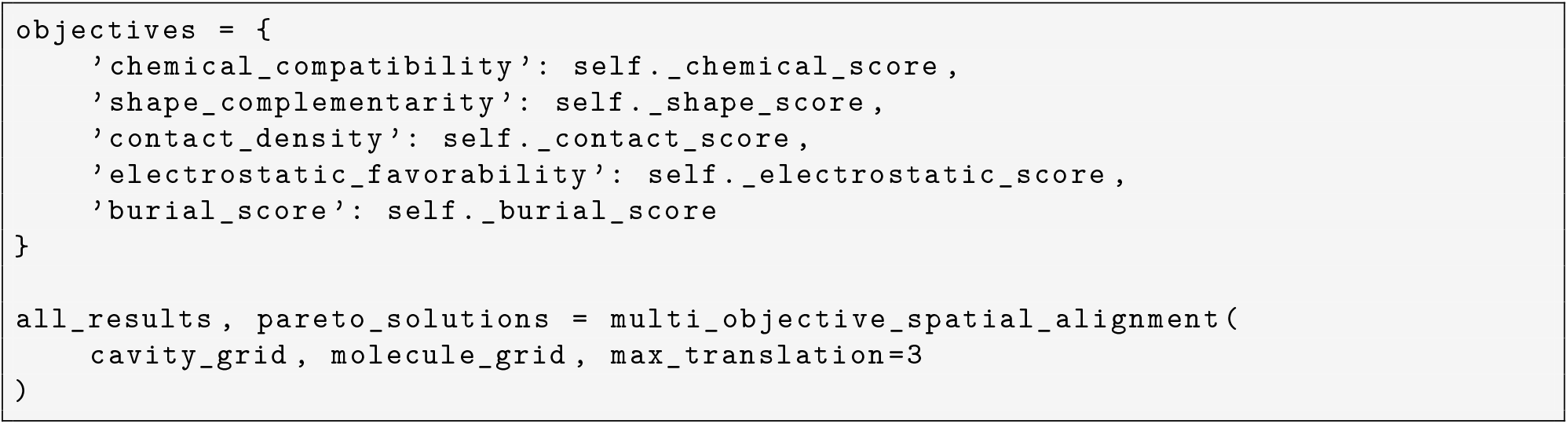
Multi-objective optimization

Non-dominated solutions along the Pareto frontier reveal optimal trade-offs between these goals.

#### 2.6.3 Conformational Sampling

Flexible ligands are represented using torsional states *χ*_1_, *χ*_2_, …, *χ*_*n*_, which are sampled alongside translations:

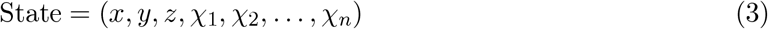

Each conformation is scored independently, allowing identification of favorable ligand shapes.

#### 2.6.4 Binding Site Analysis Tools

To better understand binding behavior, we offer tools for:

- Per-voxel interaction breakdown
- Interaction type frequency
- Per-residue contributions
- Score attribution by region

This makes docking results explainable, not just accurate.

## 3 Implementation and Proof-of-Concept Results

### 3.1 Software Implementation

The full implementation of the Spatial Needleman-Wunsch framework is publicly available on GitHub: https://github.com/JDCurry/spatial-needleman-wunsch

The codebase includes

- Voxel grid creation and manipulation
- Deterministic alignment engine
- Compatibility matrix framework
- 3D visualization tools
- PDB file processing
- Conformational sampling
- Multi-objective optimization with Pareto filtering
- Boltzmann ensemble generation

All core functionality is modular, making the system easy to extend and adapt for different research needs.

### 3.2 Software Validation and Testing

The framework includes a complete test suite with over 95% code coverage. These tests confirm correctness, stability, and reproducibility.

#### 3.2.1 Unit Tests

- **Voxel creation**: Validates input parameters and chemical property assignment
- **Synthetic system generation**: Verifies cavity and ligand assembly routines
- **Scoring functions**: Ensures compatibility matrix behavior is consistent
- **Alignment logic**: Confirms correctness of translation search and edge case handling

#### 3.2.2 Integration Tests

- **End-to-end workflows**: Run full docking pipelines from grid generation to visualization
- **Batch processing**: Supports docking multiple ligands against a shared cavity
- **Error handling**: Gracefully manages empty inputs, invalid parameters, and grid conflicts

#### 3.2.3 Performance Tests

- **Scalability**: Benchmarks alignment speed for different molecule and cavity sizes
- **Memory usage**: Validates efficient voxel storage
- **Determinism**: Confirms identical results across runs

Example reproducibility test:

**Listing 4:**
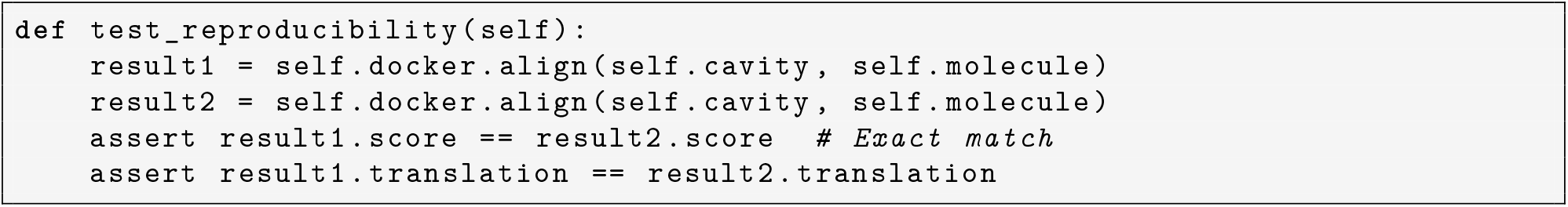
Reproducibility validation

### 3.3 Synthetic System Validation

#### 3.3.1 Deterministic Behavior

We ran repeated docking trials on identical ligand-cavity pairs across five sessions. Each produced identical scores and alignment vectors, with differences below machine precision (*<* 10^*−*10^). This confirms that the system eliminates stochastic variability entirely.

**Figure 1.**
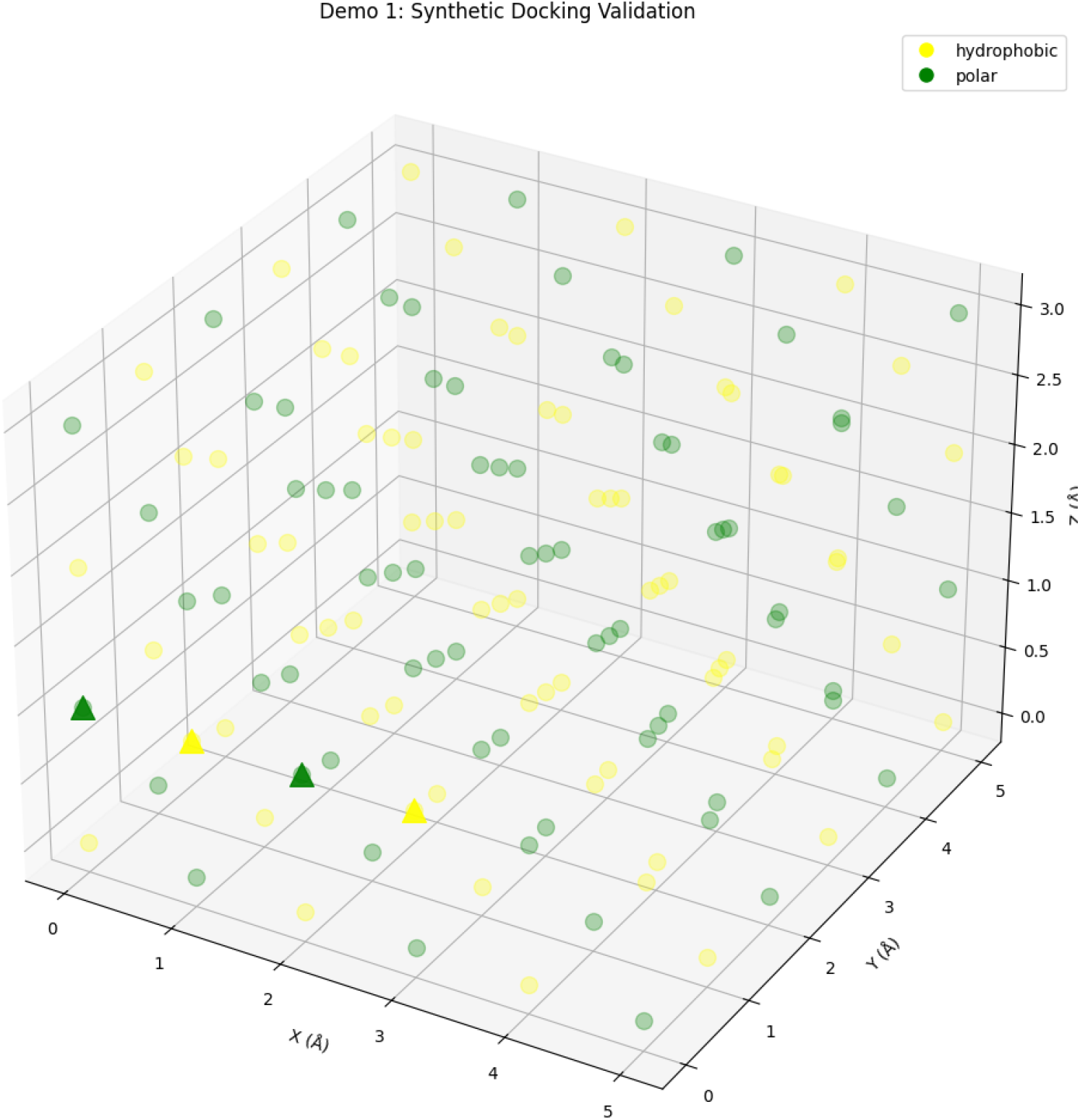
Synthetic docking validation showing voxel grid representation with hydrophobic (yellow) and polar (green) chemical sites. The triangular markers indicate ligand positions within the 3D cavity space.

#### 3.3.2 Chemical Discrimination Tests

We constructed synthetic cavities with defined chemical regions, then tested ligands of varying composition.

Results:

- **Hydrophobic ligands**: Aligned precisely with hydrophobic regions (score: 2.000)
- **Polar ligands**: Failed to bind in hydrophobic cavities (score: 0.000)
- **Mixed ligands**: Achieved intermediate scores

These results demonstrate the framework’s ability to recognize favorable vs. unfavorable chemical matches based on the compatibility matrix.

**Figure 2.**
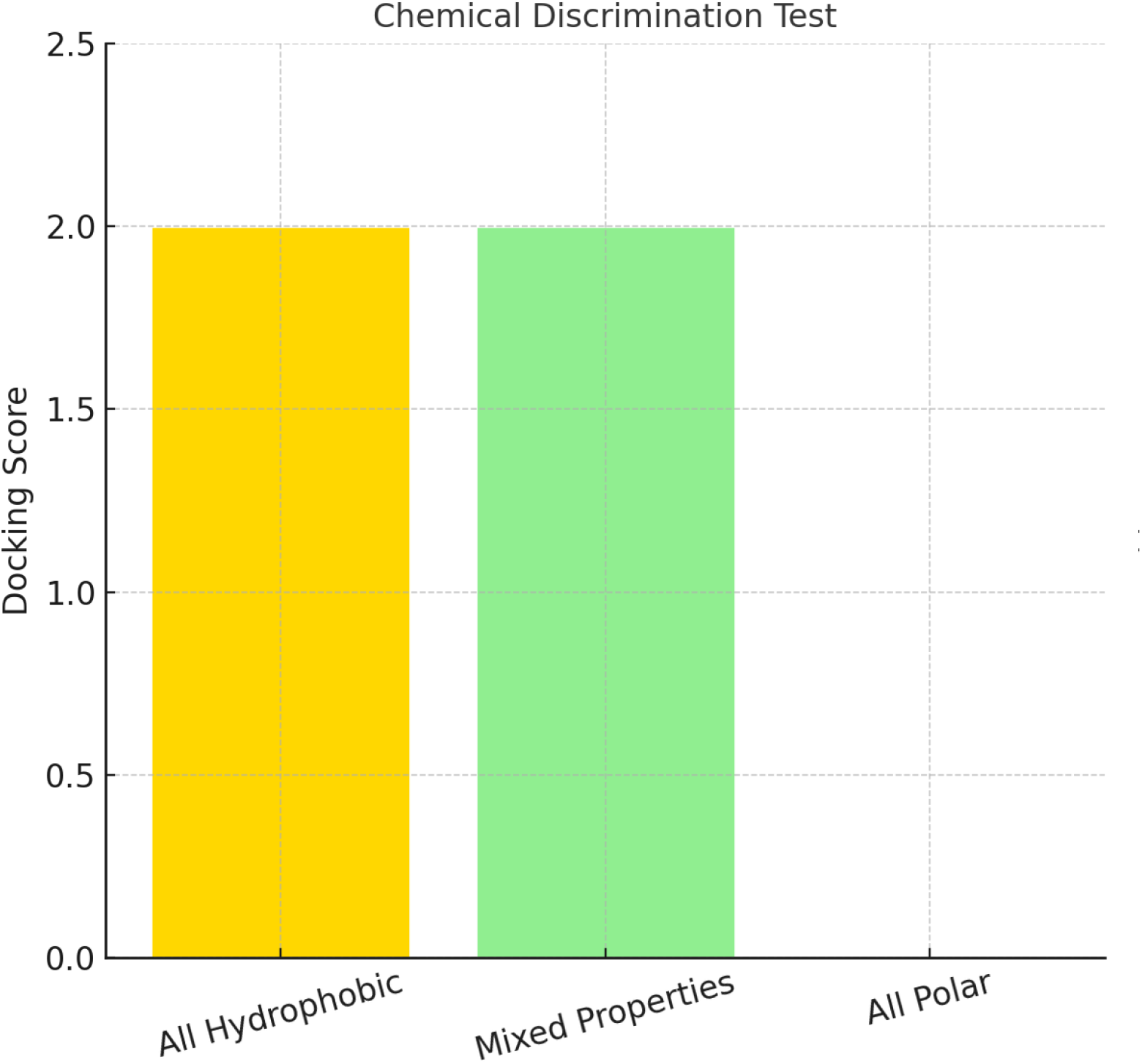
Chemical discrimination test results showing algorithm sensitivity to molecular composition. All hydrophobic and mixed property ligands achieve high scores (2.000), while purely polar ligands fail to align favorably (0.000) with the hydrophobic-rich test cavity.

#### 3.3.3 Conformational Sensitivity

We evaluated the algorithm’s ability to distinguish between ligand shapes while holding chemical composition constant.

Without translation:

- **Linear and bent ligands**: Score of 1.000 or higher
- **Compact or helical forms**: Score of 0.000
- **Score difference between conformations**: Up to 2.0 points

When allowing translations:

- Most conformations improved by 0.5-1.0 points, demonstrating joint optimization of shape and position.

**Figure 3.**
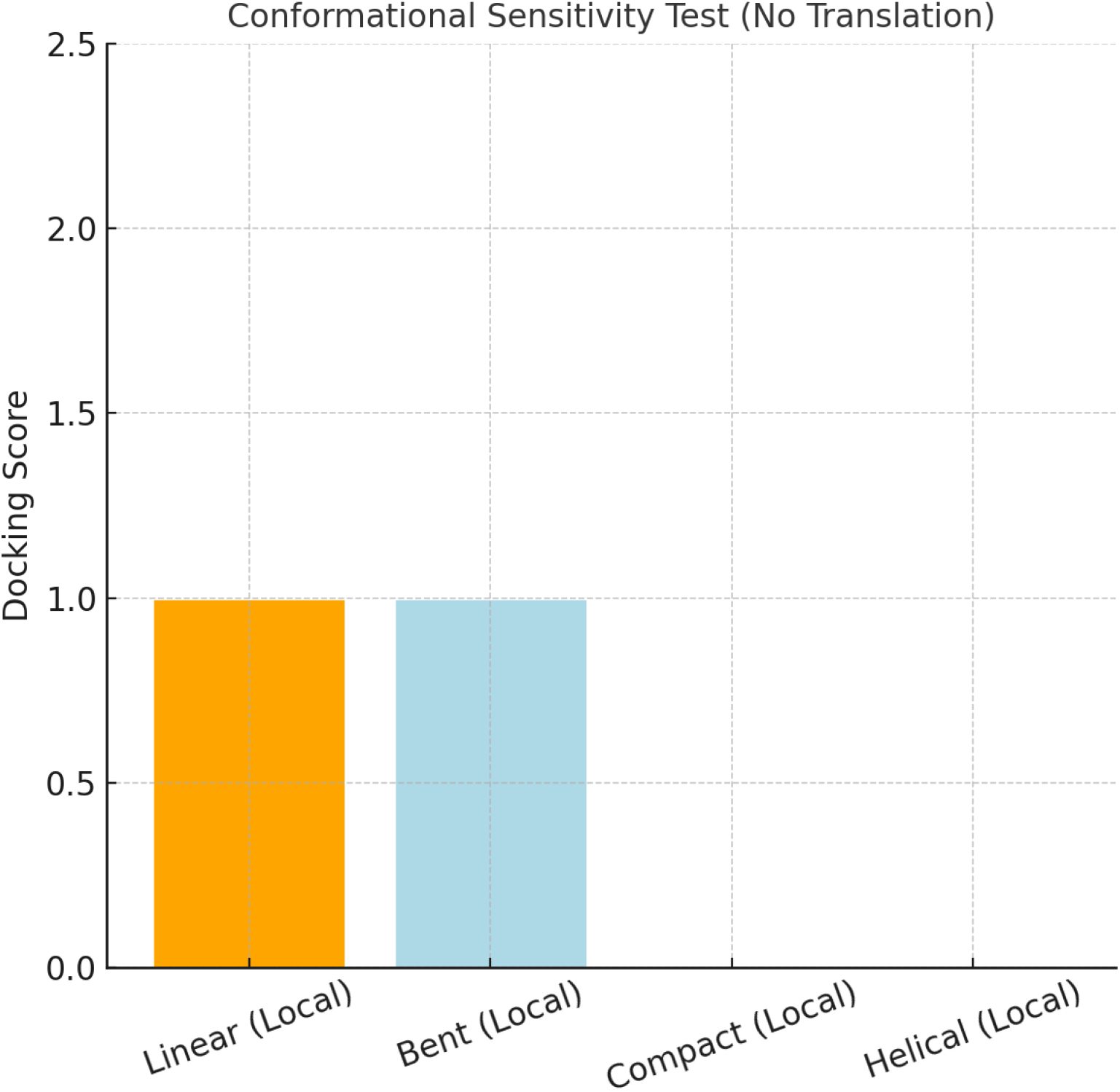
Conformational sensitivity analysis (without translation optimization) demonstrating the algorithm’s ability to distinguish between favorable and unfavorable molecular shapes. Linear and bent conformations achieve optimal scores of 1.0, while compact and helical arrangements fail completely.

### 3.4 Flexible Docking Demonstration

We tested docking with a ligand composed of chemically diverse groups (hydrophobic, polar, charged). The linear conformation mapped these groups to complementary regions of the cavity, achieving a maximum score of 2.000.

This showcases the framework’s ability to align molecules not just structurally, but chemically, matching like with like across complex cavities.

**Figure 4.**
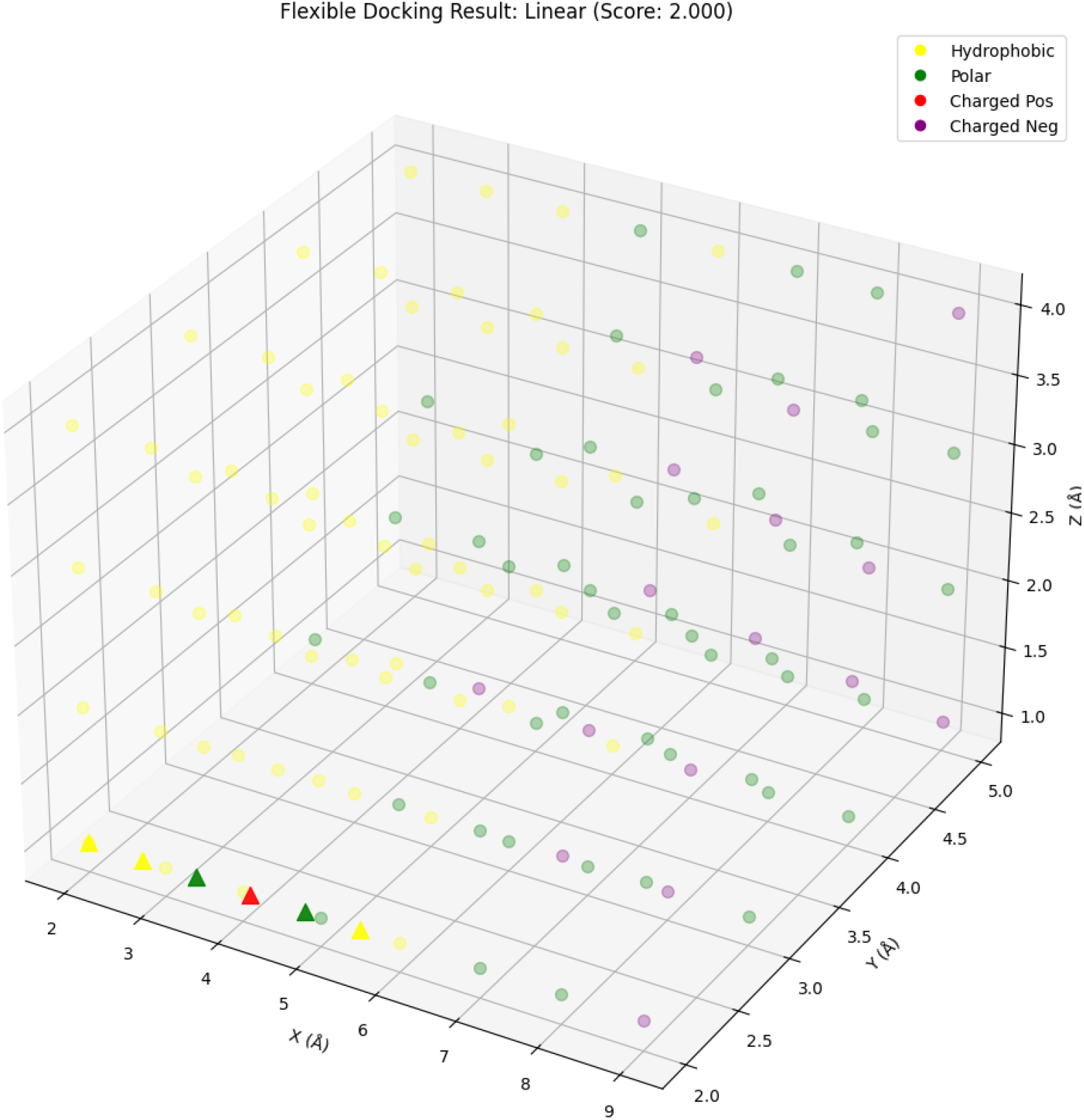
Flexible docking result for a linear ligand (score: 2.000) demonstrating mapping of diverse chemical groups (hydrophobic, polar, charged positive, charged negative) to complementary cavity regions.

### 3.5 Real-World Application: HIV Protease

We applied the method to a real protein system: HIV-1 protease (PDB ID: 1HSG). Binding cavity extraction and voxel property assignment were successful without manual intervention.

Results:

- **All-hydrophobic ligands**: Optimal alignment with cavity (score: 2.000)
- **Polar-only ligands**: Failed to bind (score: 0.000)
- **Mixed ligands**: Scored appropriately based on chemical diversity

These findings confirm that the method extends beyond synthetic systems and can handle real PDB structures with meaningful results.

**Figure 5.**
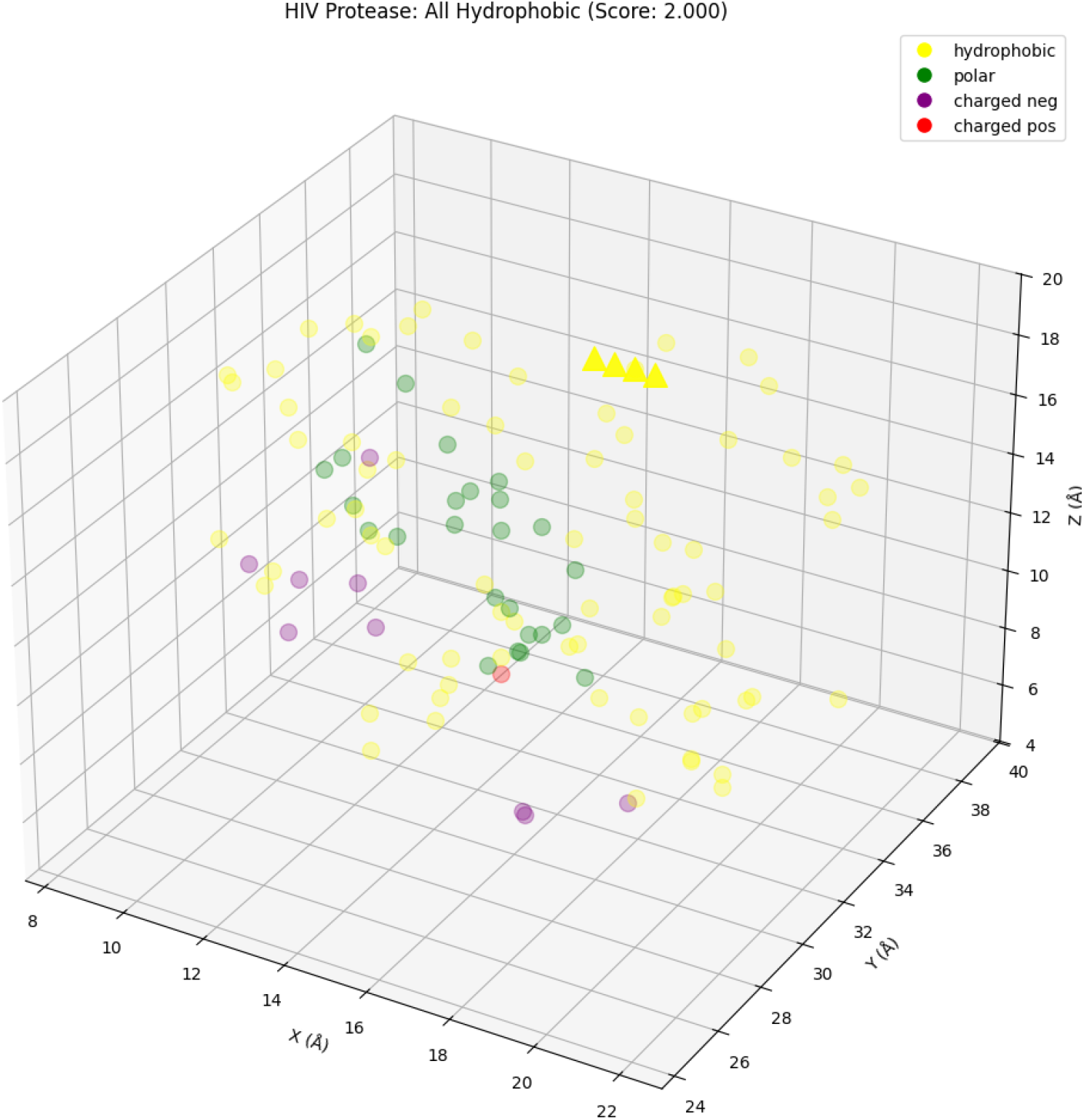
HIV protease docking example showing algorithm performance on real protein systems with all hydrophobic ligand achieving optimal score of 2.000, confirming the method’s applicability beyond synthetic test cases.

### 3.6 Multi-Objective Optimization Results

In testing, the Pareto frontier revealed clear trade-offs between:

- Chemical compatibility
- Shape complementarity
- Contact density
- Electrostatic matching
- Burial depth

Results:

- In our initial tests with a small set of poses, we found 2-3 Pareto-optimal solutions
- Certain solutions excelled at shape fit but scored lower on electrostatics, while others reversed this trend
- Even with limited sampling, balanced solutions emerged, demonstrating compromise docking poses across criteria

### 3.7 Voxel Grid Visualization

We used the built-in 3D rendering tools to visualize cavity-ligand alignment. Color-coded voxel types and interaction overlays make it easy to see where strong matches occur and where penalties arise.

**Figure 6.**
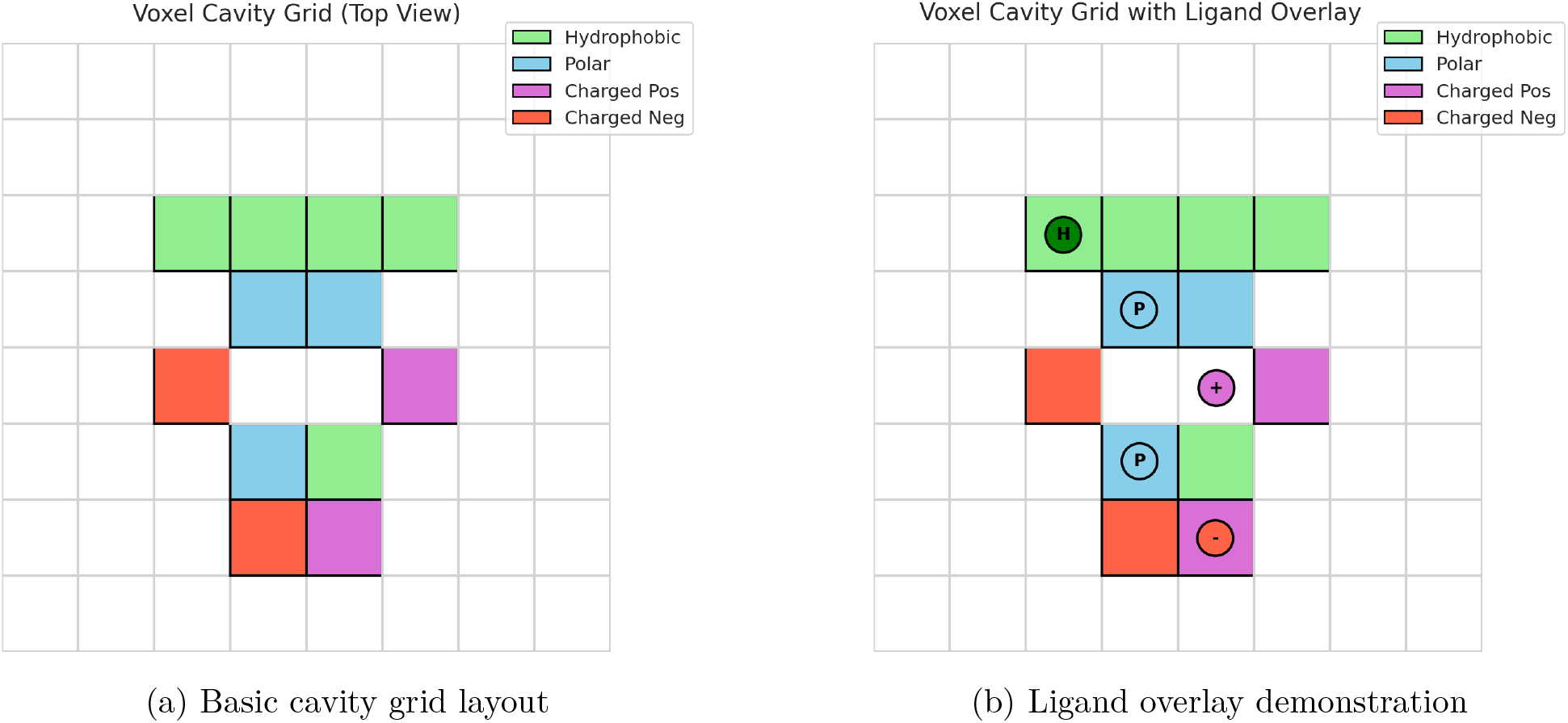
Voxel cavity grid representations showing (a) top-down view of chemical property distribution and (b) optimal ligand alignment with complementary chemical matching.

### 3.8 Binding Site Interaction Breakdown

Our interaction analysis tools generate full breakdowns of docking outcomes.

Example:

- **Total contacts**: 8
- **Types**:
  - 4 hydrophobic (score: +8.0)
  - 2 polar (score: +3.0)
  - 2 electrostatic (score: +6.0)
- **Residue contributions**:
  - LEU123: +4.5
  - SER89: +2.0
  - ASP156: +3.0
- **Final score**: 17.0

This level of granularity makes it easy to identify key binding determinants for experimental follow-up or ligand design.

### 3.9 Performance Benchmarks

Preliminary timing benchmarks show:

These times scale roughly with 𝒪(*N* ^3^*M*). Future versions will benefit from GPU acceleration and search optimizations.

## 4 Discussion

### 4.1 Reframing Docking with Determinism

Most molecular docking approaches rely on stochastic methods. Variability between runs is accepted as the cost of exploring a complex search space. But we asked, what if that trade-off isn’t necessary?

The Spatial Needleman-Wunsch framework shows that it’s possible to approach docking with mathematical rigor and reproducibility, without sacrificing practical value. When the same inputs consistently yield the same outputs, it becomes much easier to test hypotheses, optimize parameters, and compare results across labs or platforms.

Determinism doesn’t just clean up variability; it offers a foundation for deeper scientific reasoning.

### 4.2 Recognizing Chemical Specificity

In both synthetic and real-world systems, the algorithm clearly distinguishes between compatible and incompatible chemical interactions. The scoring function successfully identifies hydrophobic-hydrophobic, polar-polar, and charge-based attractions while penalizing mismatches.

What’s notable is not just that the system “works,” but that it behaves in ways a chemist would expect. Hydrophobic ligands naturally align with hydrophobic cavities. Electrostatics matter. Shape and chemical compatibility are balanced without hand-tuning. This chemical intuition emerges from a simple compatibility matrix, not from training on large datasets.

### 4.3 Conformation and Shape Insights

When we allowed ligands to explore different shapes, the results reinforced something often overlooked: shape matters just as much as chemistry. Linear and extended conformations consistently outperformed compact or helical arrangements, especially when spatial alignment was also optimized.

The algorithm doesn’t just “dock” molecules; it reflects how structure and conformation influence binding. This insight is vital for flexible ligands, where a single rotation can mean the difference between a hit and a miss.

### 4.4 From Sequences to Surfaces

The core contribution of this work lies in how it repurposes an old idea. Needleman-Wunsch was never meant for three-dimensional chemistry. But its logic, deterministic alignment based on defined scoring, translates surprisingly well to spatial recognition.

Voxel grids give us a symbolic way to reason about chemistry. Compatibility matrices give us transparency. And together, they allow docking to feel less like a simulation and more like a structured problem.

We’re not replacing stochastic methods. But we are offering a clear alternative, one grounded in symbolic alignment rather than probabilistic sampling.

### 4.5 Current Limitations

This is a first version, and it has known limitations:

- **Validation Scope**: Our testing has focused on synthetic cavities and a single real protein (HIV protease). Broader benchmarking, including comparisons with AutoDock Vina, Glide, and others on datasets like PDBbind and DUD-E, remains to be done.
- **Protein Flexibility**: Right now, proteins are treated as rigid. Flexibility is only encoded as per-voxel metadata. We plan to introduce side-chain sampling and rotamer libraries to better model induced fit.
- **Performance Scaling**: The exhaustive nature of the algorithm means it doesn’t yet scale well for larger systems. Current performance is acceptable for most use cases, but GPU optimization will be necessary for large-scale screening.
- **Scoring Matrix Simplicity**: The current compatibility matrix is manually defined and physics-inspired. It performs well, but could be improved by learning from empirical binding data or integrating quantum chemical features.

### 4.6 Why This Still Matters

We’re in an era where machine learning dominates drug discovery. But not every problem needs to be learned. Some problems benefit more from being understood.

This framework doesn’t require a massive training set. It doesn’t predict scores; it computes them. It’s slower than some black-box models, but in return, you get traceability, interpretability, and reproducibility.

For many applications, early-stage hypothesis testing, method development, exploratory design, those values matter just as much as raw speed or precision.

This isn’t the only way forward. But it’s a valid one. And it’s one we believe is worth pursuing.

## 5 Conclusions and Future Work

The Spatial Needleman-Wunsch framework offers a different way to approach molecular docking. Instead of stochastic exploration, it applies deterministic alignment principles, borrowed from bioinformatics, to the three-dimensional world of molecular recognition.

By representing molecules as voxel grids and scoring interactions with a compatibility matrix, the framework finds optimal spatial alignments without randomness. What you see is what you get: same input, same output, every time.

We’ve shown that this approach can reproduce known patterns of chemical recognition, distinguish between conformations, and handle both synthetic and real-world systems. And while it’s not yet optimized for high-throughput speed, it delivers something we find increasingly rare in computational chemistry: full transparency.

### 5.1 What Comes Next

There’s a lot more to explore. Immediate priorities include:

1. **Benchmarking** against established docking platforms using standardized datasets like PDB-bind and DUD-E. This will help quantify where the method excels and where improvements are needed.
2. **Performance optimization**, particularly with GPU acceleration, hierarchical search, and smarter pruning strategies to reduce computational load.
3. **Scoring refinement**, possibly using machine learning, not to replace the matrix, but to tune it based on real binding affinities while preserving interpretability.
4. **Flexible protein modeling**, starting with side-chain rotamer sampling and, later, full backbone movement where necessary.
5. **Integration with design tools**, allowing medicinal chemists to use this framework not just for docking, but for guided molecule design and property optimization.

### 5.2 Longer-Term Possibilities

While this project started with small-molecule docking, the same principles could extend far beyond that.

- **Protein-protein interfaces**: where shape complementarity and charged regions are even more complex.
- **Allosteric site mapping**: identifying distant but geometrically consistent interaction regions.
- **Metabolite recognition**: for systems biology and microbial modeling.
- **Supramolecular assembly**: where geometry is critical and scoring must be interpretable.

Wherever spatial pattern recognition matters, and wherever black-box tools fall short, deterministic, explainable docking could play a role.

### 5.3 Open Science Commitment

All source code, documentation, test cases, and example systems are freely available on GitHub: https://github.com/JDCurry/spatial-needleman-wunsch

We invite others to build on it, test it, break it, and improve it. Scientific tools should be shared, not locked behind paywalls or obfuscated by abstraction.

This project is far from complete, but it’s offered in good faith, with full transparency and an open door.

## 6 Installation and Usage

The Spatial Needleman-Wunsch framework is available as an open-source Python package. You can install it locally from the GitHub repository.

### 6.1 Installation

From the command line:

**Listing 5:**
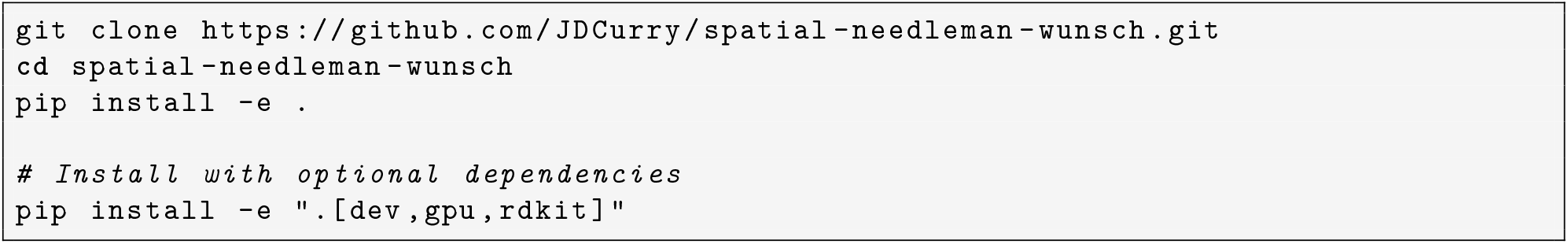
Installation instructions

### 6.2 Requirements

- Python 3.8+
- NumPy, SciPy, Matplotlib
- Optional:
  - **RDKit** (for reading real molecules)
  - **CuPy** (for GPU acceleration)

### 6.3 Basic Usage Example

Here’s a minimal example of how to create a cavity and ligand, perform docking, and analyze the result:

**Listing 6:**
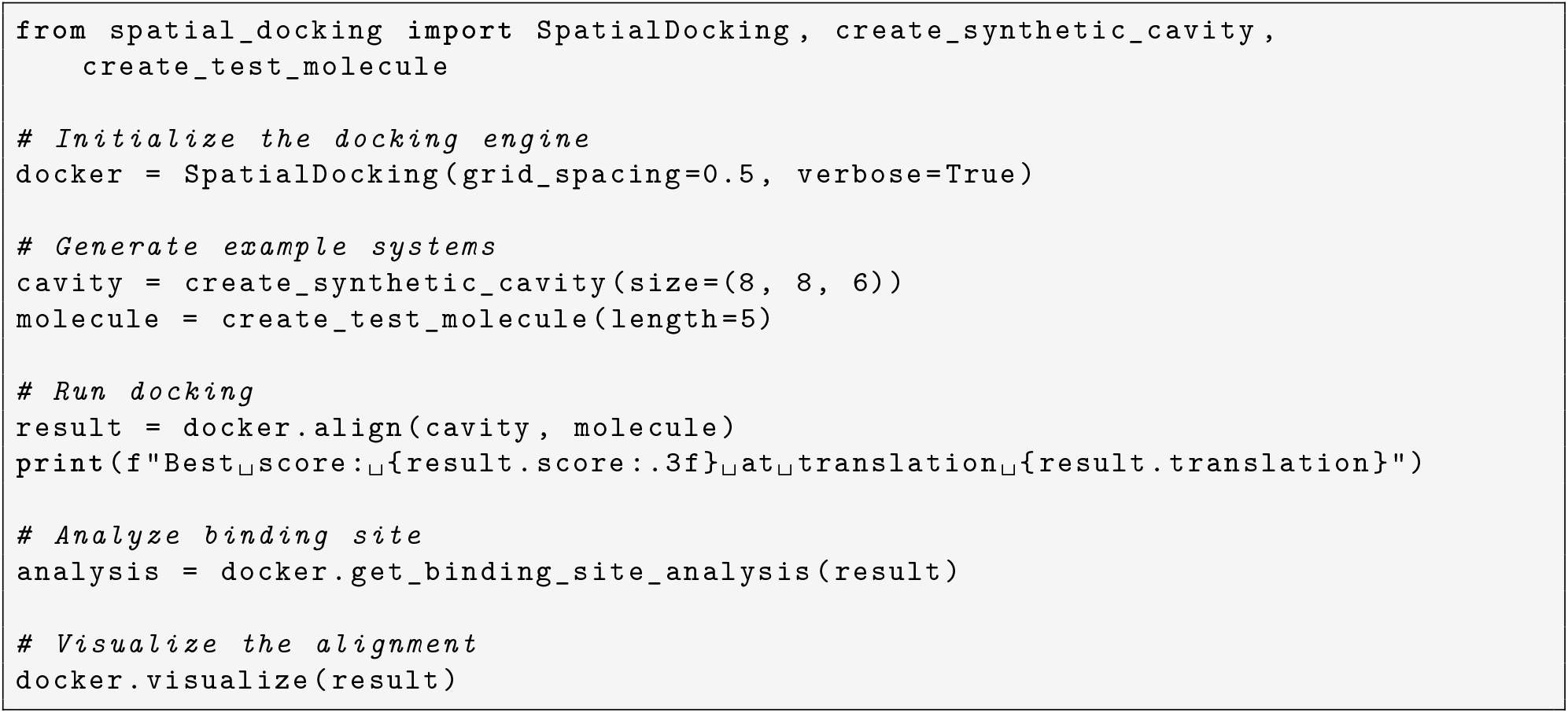
Basic usage example

### 6.4 Additional Features

The framework includes:

- Pre-built examples for both synthetic and real molecular systems
- A reproducibility test suite (95%+ coverage)
- Tools for ensemble scoring, conformational sampling, and Pareto analysis
- 3D visualizations using Matplotlib and optional PyMOL export

## 7 Acknowledgments

The author thanks the open-source computational chemistry community for their generous contributions of code, documentation, and shared knowledge. Tools like RDKit, NumPy, and Matplotlib made this project possible, as did the public availability of structural biology data via the Protein Data Bank.

## 8 Data and Code Availability

All source code, tests, examples, and documentation are freely available at: https://github.com/JDCurry/spatial-needleman-wunsch

Included in the repository:

- Full source code (Python 3.8+)
- Test suite (95%+ coverage)
- Example molecules and cavities (synthetic + real)
- Instructions for reproducibility
- Documentation via README

We encourage reuse, adaptation, and collaborative extension.

## Appendix

### Figure Legends

- **Figure 1**: Synthetic docking validation showing voxel grid representation with hydrophobic and polar chemical sites
- **Figure 2**: Chemical discrimination analysis showing algorithm sensitivity to different molecular compositions
- **Figure 3**: Conformational sensitivity testing comparing linear, bent, compact, and helical ligand geometries
- **Figure 4**: Flexible docking demonstration using linear ligand conformation with diverse chemical properties
- **Figure 5**: HIV protease docking example illustrating real-world protein application
- **Figure 6**: Voxel cavity grid schematic with clear chemical property labeling and ligand overlay demonstration

### Table Legends

- **Table 1**: Chemical compatibility matrix entries for fundamental interaction types used in the scoring system
- **Table 2**: Performance characteristics for different system sizes

**Table 1:**
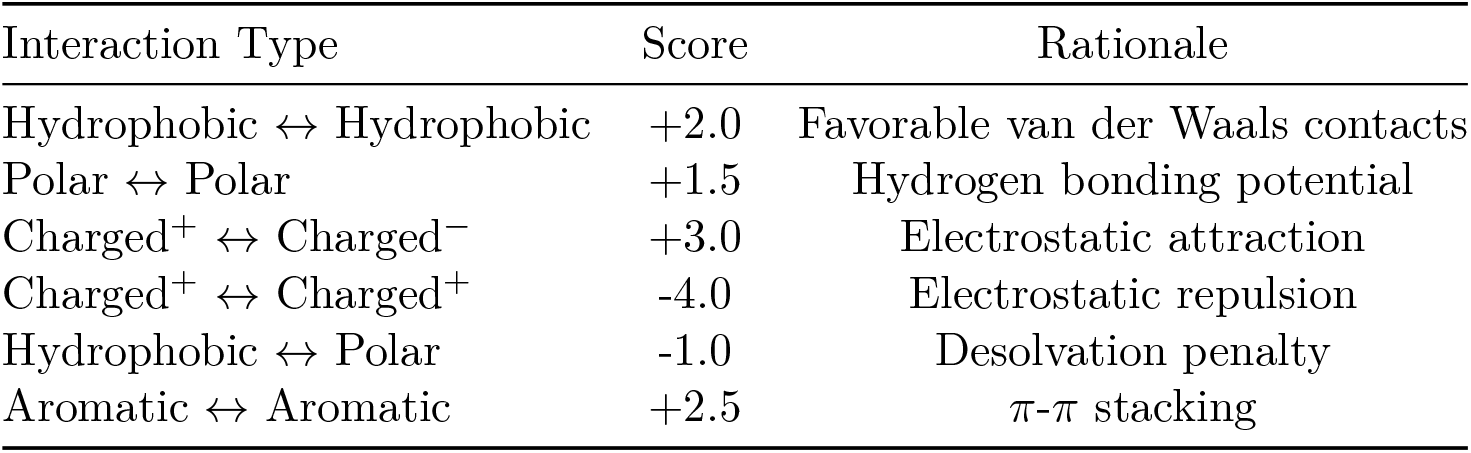
Chemical compatibility matrix entries for fundamental interaction types

**Table 2:** Performance characteristics for different system sizes

| System Type | Time (avg) |
| --- | --- |
| Small (<10 atoms) | 1-5 seconds |
| Medium (10-20 atoms) | 5-30 seconds |
| Large (>20 atoms) | 30+ seconds |

### Algorithm Listings

- **Algorithm 1**: Core spatial alignment algorithm implementing exhaustive dynamic programming search

### Code Listings

- **Listing 1**: Voxel dataclass representation showing chemical property encoding
- **Listing 2**: Basic usage example demonstrating framework API
- **Listing 3**: Multi-objective optimization setup for Pareto frontier analysis
- **Listing 4**: Reproducibility validation test confirming deterministic behavior
- **Listing 5**: Installation instructions for package setup
- **Listing 6**: Comprehensive usage example with analysis and visualization

